# Genome-wide characterization of hypothiocyanite stress response of *Escherichia coli* reveals a potential role for RclB and RclC in lowering cell envelope permeability

**DOI:** 10.1101/2024.12.06.627277

**Authors:** Julia D. Meredith, Michael J. Gray

## Abstract

Oxidative stress is one of the major methods of microbial population control and pathogen clearing by the mammalian immune system. The methods by which bacteria are able to escape damage by host-derived oxidants such as hydrogen peroxide (H_2_O_2_) and hypochlorous acid (HOCl) have been relatively well described, while other oxidants’ effect on bacteria and their genetic responses are not as well understood. Hypothiocyanite/hypothiocyanous acid (^-^OSCN/HOSCN) is one such antimicrobial oxidant. In this study, we used RNA-sequencing to characterize the global transcriptional response of *Escherichia coli* to treatment with HOSCN and observed that the response is different from the responses of *E. coli* to other oxidants such as H_2_O_2_, superoxide, or HOCl, and distinct from the responses of other bacteria such as *Streptococcus pneumoniae* and *Pseudomonas aeruginosa* to HOSCN. Furthermore, we found that deletion of the genes encoded downstream of HOSCN reductase *rclA* in *E. coli*, *rclB* and *rclC,* has a transcriptional effect on *ompC* and may play a role in membrane permeability to HOSCN.

**IMPORTANCE:** Understanding how bacteria sense and respond to oxidative stress provides insights into how our bodies interact with the microbial population within us. In this study, we have characterized the genetic response of *E. coli* to important immune oxidant hypothiocyanite and investigated the role of the *rclABC* genes in that response.

## INTRODUCTION

The innate immune system uses a variety of chemical weapons to control the population of microbes on epithelial surfaces (1). Understanding how some bacteria, whether commensal or pathogenic, are able to evade damage by the immune system while others are not, is broadly important for human health (2). Among the weapons available to the innate immune system are the production of reactive antimicrobial oxidants (3, 4). Hypothiocyanite/hypothiocyanous acid (^-^OSCN/HOSCN) is one such antimicrobial oxidant (5–7). It is related to hypohalous acids such as hypochlorous and hypobromous acid (HOCl and HOBr respectively)(8). HOSCN can be formed by three different heme peroxidase enzymes in the human body: lactoperoxidase (LPO), found secreted into multiple fluids, including saliva and breastmilk, eosinophil peroxidase (EPO), produced by eosinophils, and myeloperoxidase (MPO), which is found in leukocytes (9–12). These peroxidase enzymes catalyze the reaction between H_2_O_2_ and thiocyanate (^-^SCN) to form HOSCN (10).

HOSCN almost exclusively oxidizes thiols (13). In bacterial cells, this means that damage primarily occurs in cysteine-containing proteins, and especially glutathione and other low molecular weight thiols (14). Treatment with HOSCN causes the formation of sulfenic acids and disulfide bonds, which leads to the inhibition of many central cellular processes and potential protein aggregation (8, 12, 14).

The relationship between the immune system-produced antimicrobials and the microbiome is complex. While bacterial responses to oxidative stress is not a new field of study, there is still much unknown about how the bacteria differentially respond to the various oxidants in the human body. The transcriptional response of *E. coli* to H_2_O_2_ is different from its response to HOCl (15, 16), and the response of *Streptococcus pneumoniae* to H_2_O_2_ is different from that of *E. coli* (15, 17). While HOSCN as an antimicrobial product of LPO has been studied since as early as the 1970’s (18, 19), advances in how bacteria sense and respond to it have only come about in recent years (20–22). In fact, for a long time HOSCN was considered a highly specific antimicrobial, because mammalian cells are able to use a selenocysteine-containing thioredoxin reductase enzyme (sec-TrxR) to reduce HOSCN to its harmless precursors, SCN^-^ and water (23), while bacterial cells were thought to not possess a specific defense mechanism against it. Recently, however, we discovered that the *rcl* operon of *E. coli* encodes an efficient bacterial HOSCN reductase, which we call RclA (20), indicating that this assumption was not true and that bacteria can mount a defense against HOSCN stress.

In *E. coli*, the *rcl* operon is controlled by the transcriptional regulator RclR, which upregulates three genes in response to HOSCN: *rclA* and two additional genes called *rclB* and *rclC*, which are conserved in the *Enterobacteriaceae* family, but whose function is unknown. RclB is a small periplasmic protein, and RclC is an inner-membrane protein (6, 20). In this study, we examined the overall transcriptional response of *E. coli* to treatment with HOSCN, revealing that it is markedly different from the characterized responses of other bacteria to HOSCN, or of *E. coli* to other oxidants. We also observed that when any of the *rcl* genes are deleted, the transcriptional response of *E. coli* to HOSCN becomes more drastic, which provided us new insights into how *rclB* and *rclC* may help protect from HOSCN stress, likely by impacting the permeability of the *E. coli* cell envelope to HOSCN.

## RESULTS AND DISCUSSION

### The transcriptional response of *E. coli* to HOSCN differs from other stress responses, and from the known responses of other bacteria to HOSCN

Bacterial responses to important cellular oxidants such as H_2_O_2_ and other reactive oxygen species (ROS), have been well characterized (15, 24), as have the responses to reactive chlorine species such as HOCl (16). HOSCN reductases (RclA and its homologs Har and MerA) have recently been shown to be important in HOSCN stress resistance in *E. coli*, *Streptococcus pneumoniae*, and *Staphylococcus aureus* (20–22, 25, 26), and mutants of *E. coli* and *S. pneumoniae* lacking glutathione oxidoreductase (Gor) are very sensitive to HOSCN (21, 25, 27), but limited research has been done to explore the broader bacterial response to HOSCN or the impact of the RclA, RclB, and/or RclC proteins on that response. The entire transcriptional response to stress caused by HOSCN has only been studied in *Pseudomonas aeruginosa* strain PAO1 (28, 29), which lacks homologs of any of the *E. coli* Rcl proteins (30), and transposon sequencing has been used to identify mutants with altered HOSCN sensitivity in *S. pneumoniae* (31), which encodes only the RclA homolog Har and no homologs of RclB or RclC (21, 30).

We therefore measured the genome-wide transcriptome of *E. coli* strain MG1655 with and without treatment with 600 μM HOSCN by RNA sequencing (**FIG 1A**, **SUPPLEMENTAL DATA SET 1**). These results showed the expected strong upregulation of the *rclABC* genes (**FIG 1B**)(20). However, while the log-fold change in *rclB* and *rclC* expression was statistically significant and fairly substantial (around 3.5 fold and 4.5 fold respectively), the amount of RNA transcribed for each of these genes was much lower than that of *rclA*, despite the fact that these three genes are thought to be in an operon controlled by the RclR transcription factor acting at the *rclA* promoter (32, 33). This suggests that there is more complexity to the transcriptional regulation of the *rcl* operon than previously appreciated.

**FIG 1.**
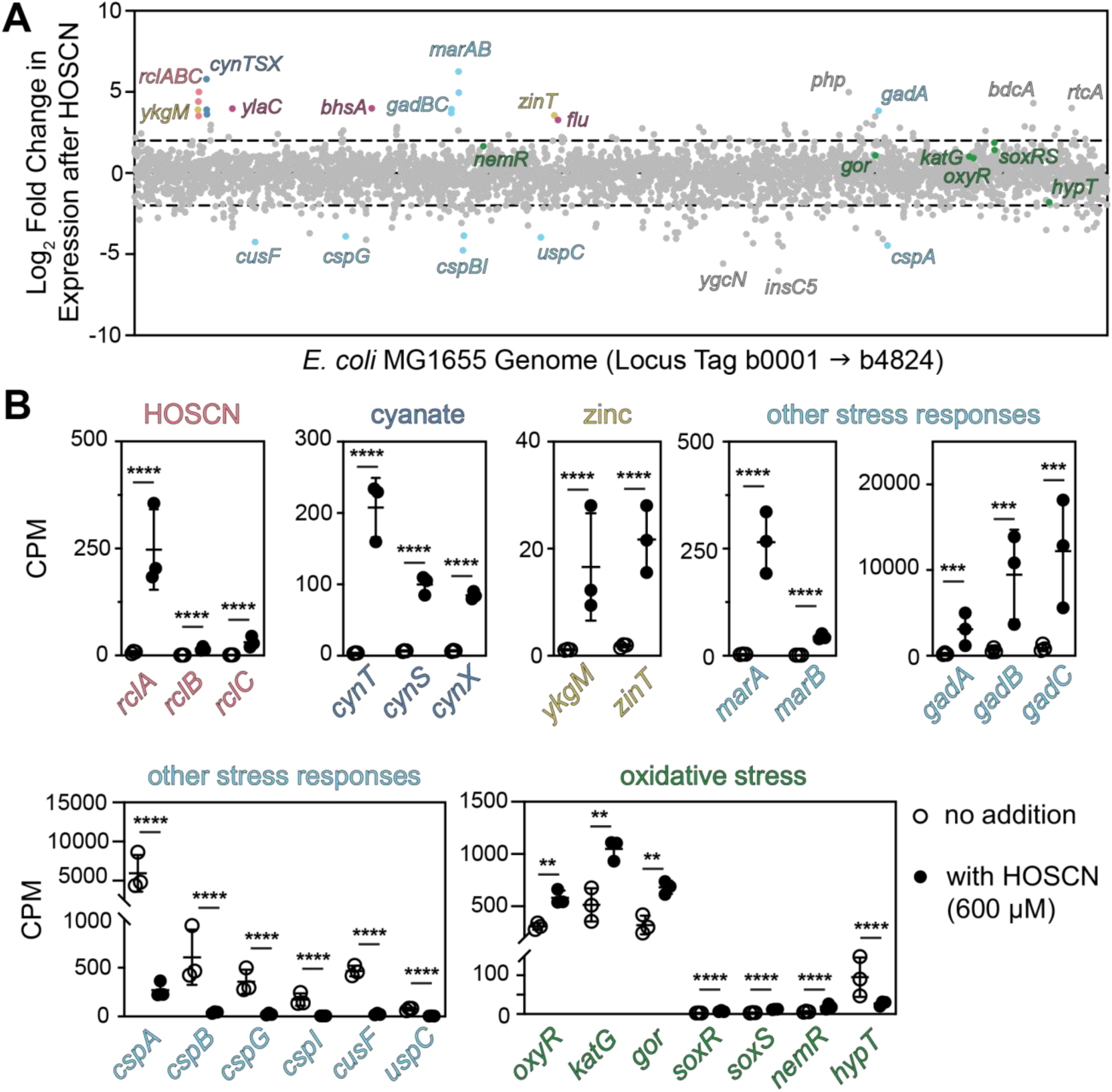
The transcriptional response to HOSCN in *E. coli*. (**A**) Transcriptomic response of *E. coli* MG1655 treated with 600 μM HOSCN compared to untreated *E. coli*. Each dot represents a gene, organized in order on the X-axis. Any gene above or below the dashed line had a log-fold change greater than 2 or -2 respectively. (**B**) The indicated genes of interest were taken from the same dataset and graphed by normalized counts per million with and without HOSCN treatment. Statistical significance calculated in Prism GraphPad by two-way ANOVA of log-transformed CPM+1 values, with significant differences in gene expression with and without HOSCN indicated: ** = P-value < 0.01, ***= P-value < 0.001, ****= P-value < 0.0001. Full RNA-seq results (**A**) are in **SUPPLEMENTAL DATA SET 1** and complete ANOVA results (**B**) are in **SUPPLEMENTAL TABLE S1**.

Other results of this experiment revealed novel aspects of HOSCN stress response in *E. coli*, which overlaps only tangentially with the well-characterized responses to other oxidants (15, 16, 24). For example, we observed significant upregulation of the *cynTSX* operon (**FIG 1B**), which encodes genes involved in transport and detoxification of cyanate (OCN^-^) (34). Expression of this operon is controlled by the OCN^-^-responsive transcription factor CynR (35), but whether this activation is due to HOSCN itself, to recognition by CynR of the SCN^-^ present in our HOSCN solution (6, 25), or to OCN^-^ contamination in our SCN^-^ solution is unclear. Results shown below (**FIGS 3** and **4**) led us to conclude that one of the second two possibilities was most likely. Genes involved in cyanate detoxification were upregulated in one study of HOSCN response in *P. aeruginosa* (28), but not the other (29).

A probably more physiologically-relevant response we observed suggested that *E. coli* experiences zinc depletion upon treatment with HOSCN (**FIG 1B**), with significant upregulation of *ykgM*, encoding a zinc-free alternative 50S ribosomal subunit protein L31B (36), and of *zinT*, encoding a periplasmic zinc-binding protein, both of which are known to be upregulated during zinc limitation (37, 38). Zinc status in *E. coli* is sensed by the transcription factor Zur, and oxidation of the zinc-binding cysteines in this protein by HOSCN could mimic the effect of low cytoplasmic zinc concentrations (39, 40). We have previously reported that RclA is sensitive to inhibition by zinc (25), and it is certainly possible that dysregulation of zinc levels has important impacts on *E. coli* survival under HOSCN stress conditions. However, *P. aeruginosa* does not show any transcriptional signatures of zinc limitation under HOSCN stress (28, 29).

Genes involved in general stress response (**FIG 1B**) that were upregulated under HOSCN treatment include the multiple antibiotic resistance genes *marA* and *marB*, which are implicated in responses to many different kinds of stress (41), as well as the acid-stress resistance genes *gadABC* (42). Some other general stress response genes were downregulated under treatment, notably including the *csp* family of genes involved in cold-shock response (43, 44), as well as *cusF*, encoding a periplasmic copper- and silver-binding protein (45), and *uspC*, encoding a so-called universal stress protein implicated in resistance to UV radiation and salt stress (46). It is not immediately clear what role any of these responses might play in HOSCN stress responses, although results shown below (**FIGS 3** and **4**) do help clarify some of those results. Again, none of these responses are mirrored in the transcriptomic results in HOSCN-stressed *P. aeruginosa* (28, 29).

Comparison of HOSCN response to other oxidative stress responses of *E. coli* revealed only limited overlaps (**FIG 1B**)(15, 16, 24). The genes encoding the H_2_O_2_-sensing transcription factor OxyR and its regulon, represented here by *katG*, encoding catalase, and *gor*, encoding glutathione oxidoreductase (24), were significantly upregulated by HOSCN, albeit by less than 2-fold, despite how important we know that Gor is to HOSCN survival in both *E. coli* and *S. pneumoniae* (25, 31). The superoxide stress response genes *soxR* and *soxS* (24) were both significantly upregulated by HOSCN, but the levels of mRNA for each of these genes was very low under both conditions.

The gene encoding the transcriptional repressor NemR, which is important to *E. coli*’s response to HOCl (47), was only slightly upregulated in response to HOSCN, while the gene encoding another transcription factor implicated in HOCl response, *hypT*, was slightly downregulated (48). In *P. aeruginosa*, Groitl *et al*. (29) observed an upregulation of *nemR* in response to HOSCN, but Farrant *et al.* (28) did not, while, in contrast, only Farrant *et al.* observed upregulation of catalase and superoxide dismutase (28). Neither study reported up- or down-regulation of *hypT* (28, 29). Consistent with the results of both *P. aeruginosa* studies (28, 29), though, we also did not observe substantial upregulation of chaperones or other heat-shock proteins, indicating that HOSCN does not cause significant amounts of protein aggregation in either species, unlike HOCl (49).

We did observe substantial HOSCN-dependent upregulation of *bhsA*, also known as *comC* (**FIG 1B**), which encodes a small periplasmic / outer membrane protein that impacts biofilm formation and appears to modify the permeability of the outer membrane to copper by an unknown mechanism, playing a role in increasing resistance to copper toxicity (50, 51). We will return to this gene in the sections below (**FIGS 3, 4**, and **6**) for a more thorough discussion of the implications of this result.

Finally, a variety of other genes with poorly characterized functions were notably up- or down-regulated as well, but what role these genes might play in HOSCN response is not clear. These included the genes encoding the putative inner membrane protein YlaC (52), the putative zinc-dependent hydrolase Php (53), the c-di-GMP binding protein BdcA (54), the RNA 3’-terminal phosphate cyclase RtcA (55), and the putative oxidoreductase YgcN (52).

### Knockouts of *rcl* genes are more sensitive to HOSCN than wild-type *E. coli*

The *rclABC* operon of *E. coli* is of particular interest to our group (20, 25, 30, 33). The functions of RclB and RclC, encoded in an operon downstream of *rclA* and mostly conserved only among members of the Enterobacteriacea (30, 33), have not been elucidated. RclB is a predicted periplasmic protein belonging to the DUF1471 family of protein domains of unknown function (56, 57) and RclC is a predicted inner membrane protein (33) of the DUF417 family (56), but little else is known about their functions. It is important to note that neither RclB nor RclC possess cysteine residues, and therefore are unlikely to be oxidized directly by HOSCN (13). An RclC homolog called RcrB which is found in uropathogenic *E. coli* plays an important and specific role in HOCl resistance, but its mechanism of action has also not been characterized (58). Null mutations in *rclB* or *rclC* confer substantial sensitivity to HOSCN, comparable to the very high sensitivity of an Δ*rclABC* triple mutant (25) at high HOSCN concentrations (**FIG 2, SUPPLEMENTAL FIG S1**). Notably, the sensitivities of Δ*rclB* and Δ*rclC* mutants were very similar to one another under all tested conditions, suggesting that they may function together.

**FIG 2.**
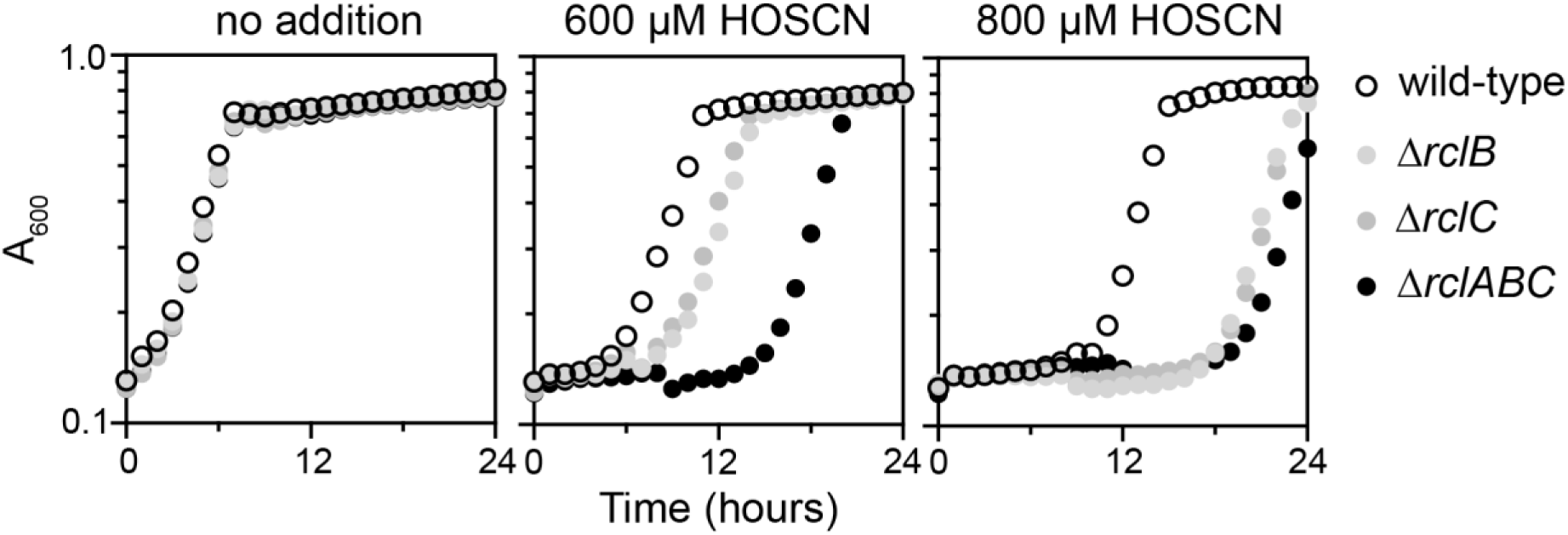
RclB and RclC are required for HOSCN stress resistance in *E. coli*. *E. coli* MG1655 wild-type, MJG0047 (MG1655 Δ*rclB*), MJG0013 (MG1655 Δ*rclC*::*cat*^+^), and MJG0901 (MG1655 Δ*rclABC*) were inoculated into M9 minimal medium containing the indicated concentrations of HOSCN and incubated at 37°C with shaking for 24 hours in a Tecan Sunrise plate reader, measuring A_600_ every 30 minutes. Each graph shows the mean of four technical replicates, and two additional experimental replicates are shown in **SUPPLEMENTAL FIG S1**.

### *E. coli* mounts a specific response to HOSCN when *rcl* genes are mutated

*E. coli*’s response to HOSCN was very noticeably different when any of the *rcl* genes are mutated when compared to the wild-type (**FIG 3**). However, surprisingly, the response between individual mutations was not extremely variable, suggesting that RclA, RclB, and RclC form a coordinated defense against HOSCN and that the loss of any one of these proteins leads to similar breakdowns in the ability of the cell to respond appropriately to HOSCN stress, at least at a transcriptional level. Across the board, we observed stronger upregulation of genes involved in metal stress response, envelope stress response, methionine synthesis, and oxidative stress response when any of the *rcl* genes was disrupted. In **FIG 4**, we present a set of representative genes of interest in more detail to compare induction by HOSCN in the wild-type and each mutant strain.

**FIG 3.**
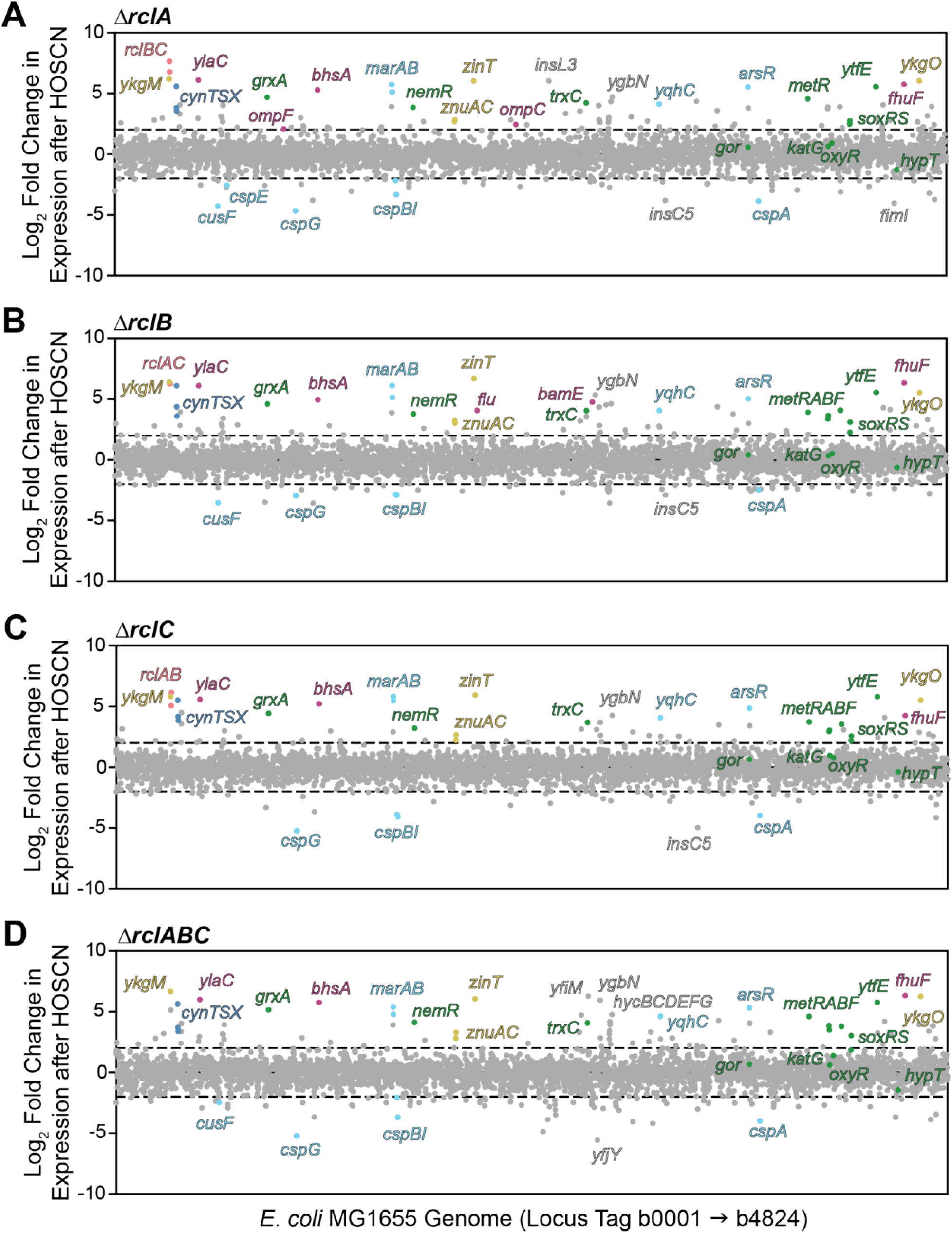
The transcriptional response of *E. coli* Δ*rclA*, Δ*rclB*, Δ*rclC*, and Δ*rclABC* mutants to HOSCN. *E. coli* transcriptional response to 600 μM HOSCN in strains (**A**) MJG1958 (MG1655 Δ*rclA*), (**B**) MJG0047 (MG1655 Δ*rclB*), (**C**) MJG0013 (MG1655 Δ*rclC*::*cat*+), or (**D**) MJG0901 (MG1655 Δ*rclABC*). Each dot represents a gene, organized in order on the X-axis. Any gene above or below the dashed line had a log-fold change greater than 2 or -2 respectively. Full RNA-seq results are in **SUPPLEMENTAL DATA SET 1**.

**FIG 4.**
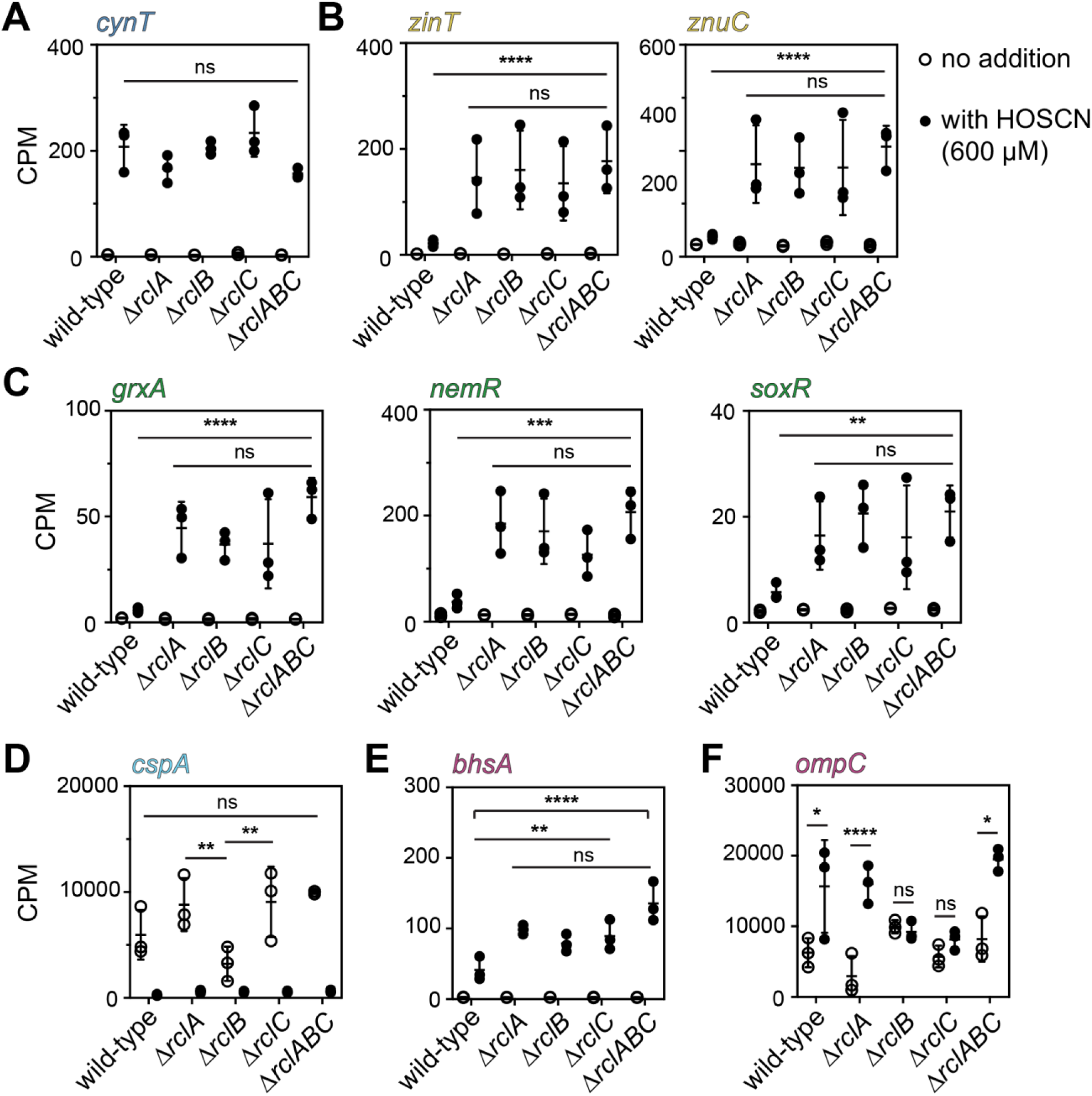
Expression of selected genes, illustrating notable features of the transcriptional response of *E. coli* Δ*rclA*, Δ*rclB*, Δ*rclC*, and Δ*rclABC* mutants to HOSCN. CPM values with and without 600 µM HOSCN treatment (see **FIGS 1**, **3**) for (**A**) the cyanate response gene *cynT*, (**B**) the zinc starvation response genes *zinT* and *znuC*, (**C**) the redox stress response genes *grxA, nemR*, and *soxR*, (**D**) the cold shock gene *cspA,* (**E**) the copper resistance gene *bhsA*, and (**F**) the *ompC* outer membrane porin gene in strains MG1655, MJG1958 (MG1655 Δ*rclA*), MJG0047 (MG1655 Δ*rclB*), MJG0013 (MG1655 Δ*rclC*::*cat*+), and MJG0901 (MG1655 Δ*rclABC*). Statistical significance calculated in Prism GraphPad by two-way ANOVA of log-transformed CPM+1 values, with significant differences of interest indicated: ns = not significant, * = P-value < 0.05, ** = P-value < 0.01, ***= P-value < 0.001, ****= P-value < 0.0001. Note that panel (**F**) is the only one in this figure that shows comparisons between expression with and without HOSCN, all of which were significant for every strain in all of the other panels. Full ANOVA results in **SUPPLEMENTAL TABLE S1**.

Many of the general stress response genes that were upregulated in the general response of *E. coli* were equally upregulated in the *rcl* mutant strains, including *cynT* (**FIG 4A**), indicating an equal OCN^-^ stress response that was not dependent on the *rcl* operon. This reinforced our conclusion that expression of *cynTSX* was not likely to be a *bona fide* HOSCN response. Similarly, for example, the *marA* and *marB* genes, encoding regulators of genes involved in responses to a variety of toxic compounds (41), notably those encoding the AcrAB efflux pump (59), were strongly upregulated by HOSCN regardless of the presence of *rclABC* (**FIGS 1**, **3**, **SUPPLEMENTAL DATA SET 1**). However, *acrA* and *acrB* were not substantially upregulated in any HOSCN-treated strain (**SUPPLEMENTAL DATA SET 1**), making the significance of *marAB* upregulation unclear.

More commonly, we observed stress response genes that were much more strongly upregulated in all four *rcl* mutant strains than in the wild-type, but which were not significantly different among the mutant strains (**FIG 4B**, **C**). For example, the zinc starvation response, which was seen in wild-type (**FIG 1B**), is much more evident under mutated *rcl* conditions (**FIG 4B**), as demonstrated by the greater upregulation of *zinT* and the strong upregulation of *znuC*, which encodes part of an ABC transporter which is involved in zinc uptake (38, 60). There is also a much more prominent signature of oxidative stress response to HOSCN in the *rcl* mutant strains. For example, expression of *grxA,* encoding one of the four glutaredoxins found in *E. coli* (61), is raised in the *rcl* mutants (**FIG 4C**), which was somewhat surprising, considering that we have recently reported that mutants of *E. coli* lacking *grxA* are not more sensitive to HOSCN than the wild-type (25). Because of HOSCN’s ability to efficiently oxidize low molecular weight thiols (13, 62, 63), the connection between glutathione and bacterial resistance to HOSCN has recently been of interest in the field. *E. coli* Δ*gor* mutants (lacking glutathione oxidoreductase)(52, 64), for instance, are highly sensitive to HOSCN (25). In *S. pneumoniae*, double mutants of *gor* and *har* (encoding the RclA homolog of that species)(21) are also highly sensitive to HOSCN (21, 27), and indeed we did see a general upregulation of *gor* in *E. coli* under HOSCN treatment, however this was consistently 2-fold induction or less and was not substantially affected by mutation of the *rcl* genes (**FIGS 1**, **4**, **SUPPLEMENTAL DATA SET 1**).

When *E. coli* is exposed to HOCl stress, there is strong upregulation of the redox sensitive transcription factor-encoding gene *nemR* (47). Under HOSCN stress, we observed only a modest upregulation of *nemR* in the wild-type (**FIG 1B**), with a much stronger upregulation when any or all of the three *rcl* genes are missing (**FIG 4C**).

Likewise, *soxR,* encoding a transcription factor that responds to superoxide, redox-cycling drugs, and nitric oxide (41, 65), is expressed in much higher amounts in *rcl* mutants (**FIG 4C**). NemR is a bleach-sensing transcriptional regulator that responds to HOCl stress through oxidation of its cysteine residues (47, 66), and SoxR senses various oxidative stresses via an oxidation-sensitive cysteine-based iron-sulfur cluster (65). Logically, it makes sense that there would be more protein cysteine oxidation in a strain of *E. coli* missing specific defenses against HOSCN (13, 62, 63). The upregulation of the MetJ regulon (67) in *rcl* mutants (**FIG 3**, **SUPPLEMENTAL DATA SET 1**) suggests that methionine may also be oxidized under these conditions, although this was unexpected since HOSCN does not efficiently react with methionine *in vitro* (13). It is interesting, however, that across the board there is little difference in the amount of upregulation of these oxidative stress responses in individual mutants.

Mutants lacking *rclB* or *rclC* are not as sensitive to HOSCN as a mutant of *rclA* (20) or as the triple knockout (**FIG 2**), and *E. coli* doesn’t make as much *rclB* or *rclC* as *rclA* (**FIG 1B**), but the different mutant cells seem to be perceiving the same amount of oxidative stress, based on their transcriptional responses.

The impact of *rcl* mutations on the down-regulated *csp* genes (**FIG 1B**) was less consistent. For example, while there was no significant difference in *cspA* expression between wild-type and any of the mutant strains (**FIG 4D**), expression of that gene was lower in the absence of HOSCN in the Δ*rclB* mutant than in either the Δ*rclA* or Δ*rclC* mutants. The mechanisms and functions of the proteins encoded by the *csp* genes are not generally well-understood (44), and the physiological relevance of this observation, if any, is unclear.

Perhaps the most interesting to us was the very small set of genes whose expression did differ among the various *rcl* mutant strains. Two that stood out were *bhsA* and *ompC* (**FIG 4E**, **F**). The *bhsA* gene encodes a small periplasmic protein of the same DUF1471 family as RclB, has been shown to affect both biofilm formation and cell envelope permeability to copper (50, 51), and was the only gene regulated by treatment with HOSCN that was more significantly upregulated in the triple knockout of *rclABC* than in the single mutants (**FIG 4E**, **SUPPLEMENTAL DATA SET 1**). This already suggested to us that there might be a link between cell envelope permeability, HOSCN treatment, and the function(s) of the *rcl* operon. Meanwhile, *ompC* encodes one of the major outer membrane porins (OMPs) of *E. coli* (68), and had a very striking and unique pattern of expression. Transcription of *ompC* was significantly induced by HOSCN in wild-type, Δ*rclA*, and Δ*rclABC* strains, but not in either of the Δ*rclB* or Δ*rclC* mutants (**FIG 4F**, **SUPPLEMENTAL DATA SET 1**). The genetic regulation of the OMPs is complicated and affected by over a dozen known transcriptional regulators (69), but in general, *ompC* transcription increases in the presence of stressors affecting the cell envelope. This therefore suggested a model in which HOSCN might accumulate in the periplasmic space of wild-type or Δ*rclA* mutant strains (damaging the cell envelope and inducing *ompC* expression) but is freely permeable through the cell envelope of mutants lacking RclB or RclC. In the presence of cytoplasmic RclA (20), that HOSCN would be rapidly degraded (preventing envelope damage and *ompC* induction). However, according to this model, in the mutant lacking all three Rcl proteins, HOSCN would be relatively stable and be able to react with and damage the cell envelope, restoring *ompC* induction in the Δ*rclABC* strain (**FIG 4F**). We therefore turned our attention to the potential role of outer membrane and cell envelope permeability in *E. coli* resistance to HOSCN.

### Mutations in outer membrane porins protect *E. coli* from HOSCN

OmpC, OmpF, and OmpA are the major OMPs in *E. coli* (68, 69). The loss of outer membrane permeability via OMP regulation has often been associated with antibiotic resistance in diverse bacteria (70), and deletions of either *ompC* or *ompF* are reported to protect *E. coli* against the HOSCN-generating enzyme LPO (71). We therefore made deletions of both *ompC* and *ompF* in our *E. coli* strain background to test those mutants’ sensitivity to HOSCN treatment (**FIG 5**, **SUPPLEMENTAL FIG S2**). Single mutants lacking either *ompC* or *ompF* did not have any substantial sensitivity or resistance to HOSCN and grew equally well to the wild-type at moderate HOSCN concentrations (400 or 600 µM). Both Δ*ompC* and Δ*ompF* mutants were slightly more resistant to 800 µM HOSCN than the wild-type strain (**SUPPLEMENTAL FIG S2**). As expected due to the loss of most of the porins allowing solute transfer across the outer membrane (68), the Δ*ompCF* double knockout grew more slowly without stress than the other strains but it also displayed strong resistance to HOSCN. At higher HOSCN concentrations (600 or 800 µM), the double knockout strain grew better than the wild-type or the single knockouts (**FIG 5**, **SUPPLEMENTAL FIG S2**). These results confirmed that reduction of outer membrane permeability by disrupting OMP function can confer HOSCN resistance in *E. coli*.

**FIG 5.**
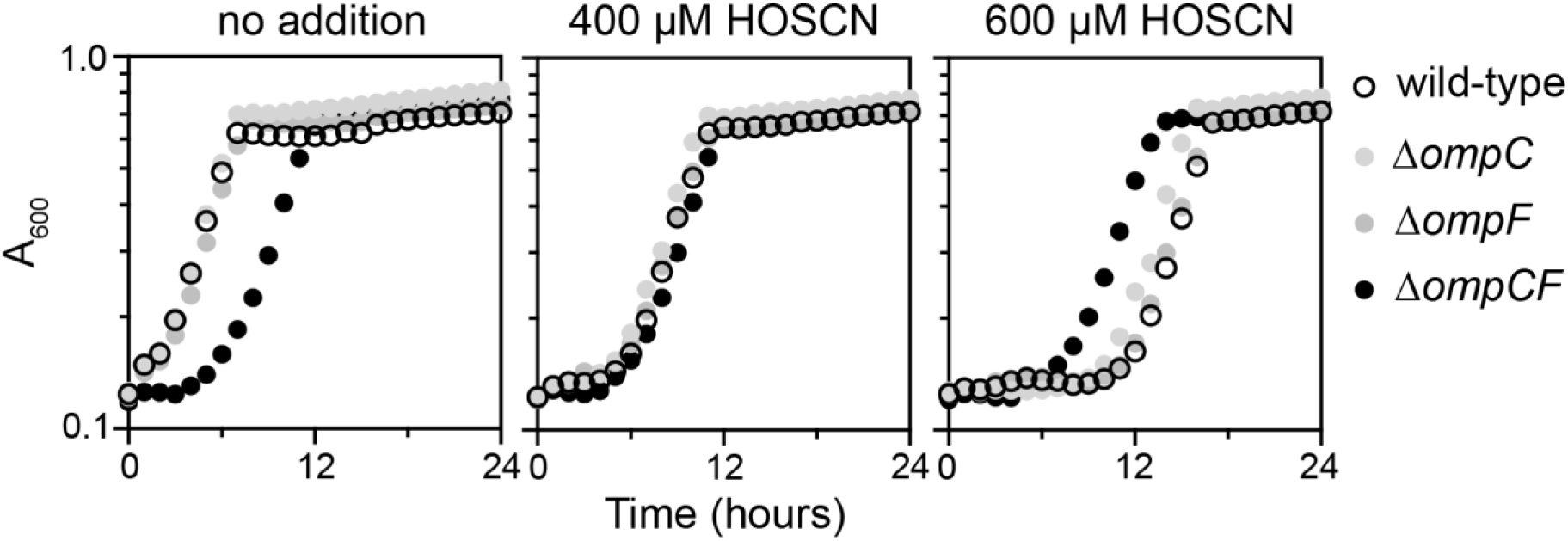
Mutations in *ompC* and *ompF* protect *E. coli* against HOSCN stress. *E. coli* MG1655 wild-type, MJG2411 (MG1655 Δ*ompC*), MJG2412 (MG1655 Δ*ompF*), and MJG2429 (MG1655 Δ*ompC* Δ*ompF*::*kan*^+^) were inoculated into M9 minimal medium containing the indicated concentrations of HOSCN and incubated at 37°C with shaking for 24 hours in a Tecan Sunrise plate reader, measuring A_600_ every 30 minutes. Each graph shows the mean of four technical replicates, and two additional experimental replicates are shown in **SUPPLEMENTAL FIG S2**.

### RclB is homologous to other small periplasmic stress response proteins

RclB belongs to a family of small proteins called the DUF1471 family, which are small periplasmic proteins exclusive to the *Enterobacteriaceae*, only a few of which have any functional annotation (57). *E. coli* MG1655 encodes 10 DUF1471 paralogs (**SUPPLEMENTAL TABLE S2**)(52). The amino acid sequences of the most similar paralogs to RclB are shown in **FIG 6**. Most of these have no known functions, but notable among them are BhsA, discussed above (**FIG 4E**), and YjfN, which uses a hydrophobic alanine on its C-terminus to interact with the protease DegP, directing the degradation of OmpA under certain envelope stress conditions (72). BhsA is known to be associated with the outer membrane, where it reduces the permeability of the cell envelope to copper and therefore increases the resistance of *E. coli* to copper stress (50), but the exact mechanism by which it affects permeability is not known. Some of the other DUF1471 proteins are also associated with specific stress responses (*e.g.* YhcN with oxidative stress (73), McbA with biofilm formation (74), or YahO with radiation resistance (75)), but nothing is known about how they contribute to these responses.

**FIG 6.**
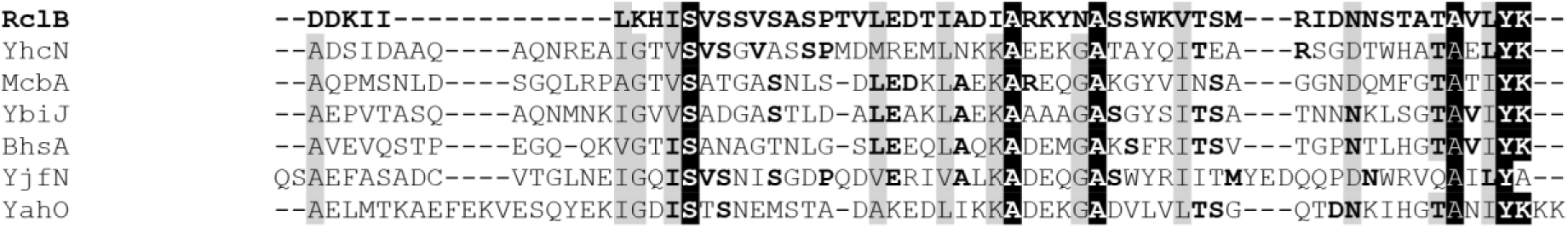
Alignment of RclB with other *E. coli* DUF1471 proteins. The EMBL MUSCLE multiple alignment tool (89) was used to compare RclB to the six most similar of the other DUF1471 proteins in *E. coli* MG1655 (52)(see **SUPPLEMENTAL TABLE S2** for additional information about these and the other five DUF1471 domains encoded by *E. coli* MG1655).

A separate search for structural homologs of RclB, using the COFACTOR tool (76), identified the *E. coli* RcsF protein as containing a highly similar three-dimensional structure (RMSD^a^ = 1.47 Å)(**FIG 7**). RcsF is a sensor lipoprotein that forms a complex with OmpC, OmpF, or OmpA (called the RcsF-OMP complex), inserting into the β-barrel pores of those OMPs and using its surface-exposed lipidated loop to detect membrane damage (77). RcsF is required for activation of the envelope stress-responsive Rcs system (77). While RclB is not very similar to RcsF at the primary sequence level (18% identical, 30% similar; **SUPPLEMENTAL TABLE S2**) and does not possess the loop which forms the lipidated section of RcsF (**FIG 7C**, lower left corner), the structural homology could still indicate some sort of interaction with OMPs, and indeed, based on the results we show here, we hypothesize that interaction with or regulation of OMPs to control outer membrane permeability might be a common function for all of the stress-responsive DUF1471 proteins (**FIG 6**, **SUPPLEMENTAL TABLE S2**).

**FIG 7.**
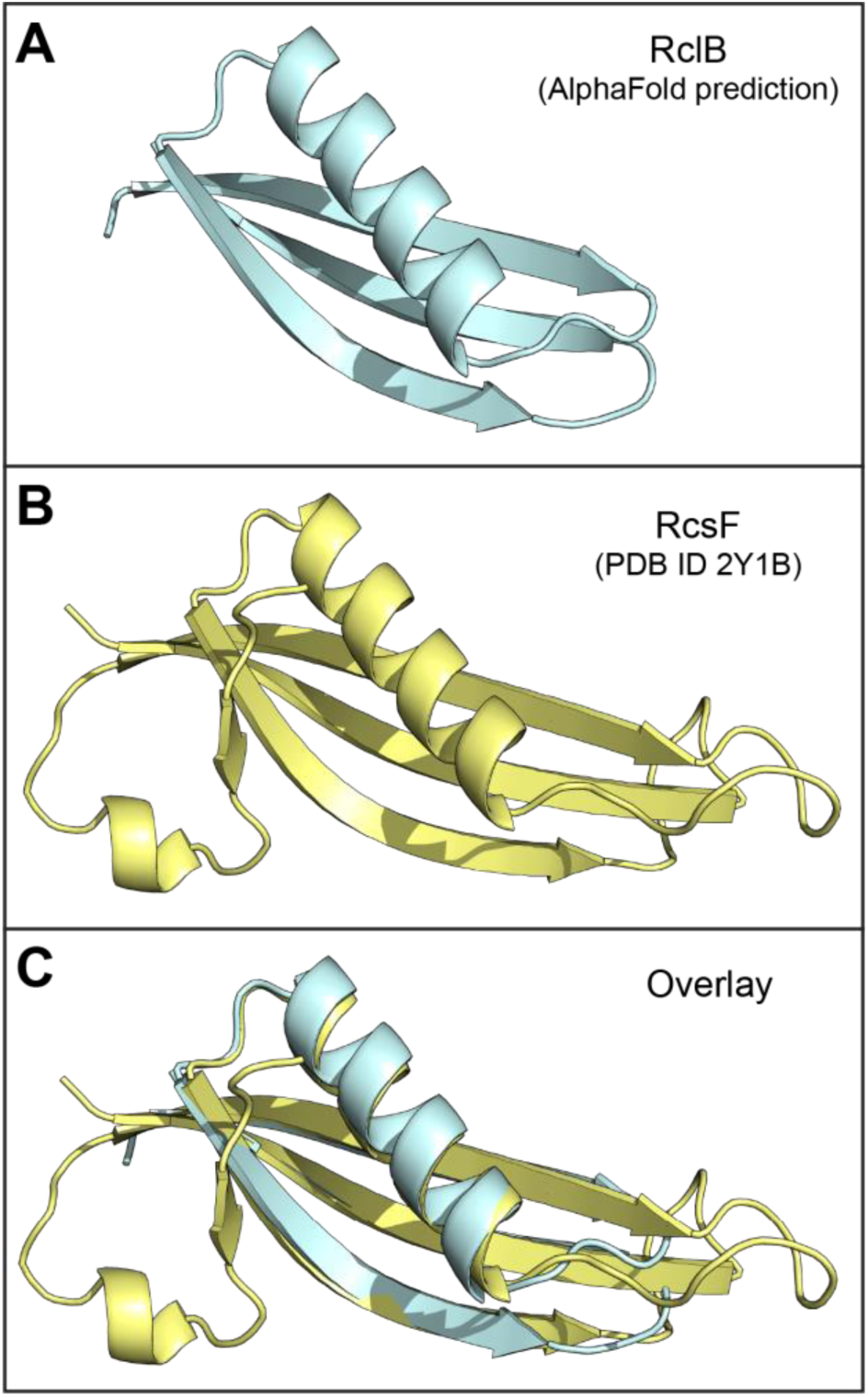
RclB is a structural homolog of the OMP-interacting lipoprotein RcsF. (**A**) The AlphaFold prediction of the structure of RclB (85, 86) and (**B**) the crystal structure of RcsF (90) were identified as structural homologs (**C**) using COFACTOR (76), which calculated a RMSD^a^ score = 1.47 Å. Protein structures were visualized using Pymol.

### Neither HOSCN treatment nor mutation of *rclB* and *rclC* has any effect on abundance of OMPs

Since Δ*ompCF* mutants are resistant to HOSCN (**FIG 5**) and RclB is homologous to two proteins known to interact with or regulate OMPs at the post-transcriptional level (*i.e.* YjfN and RcsF), we tested whether either exposure to HOSCN or *rcl* mutations affected the abundance or ratio of the OmpC, OmpF, or OmpA proteins in *E. coli* (68, 69). Under our growth conditions, OmpC, OmpF, and OmpA were all highly abundant (**FIG 8A**, **SUPPLEMENTAL FIG S3**). We were unable to consistently isolate high quality outer-membrane fractions from *E. coli* exposed to high HOSCN concentrations, but exposure to 200 µM HOSCN (sufficient to strongly induce transcription of the *rclABC* operon)(20) had no significant effect on the abundance of any of the OMPs (**FIG 8B**). The Δ*rclABC* mutant had slightly less OmpA than the wild-type under both non-stress and HOSCN treatment conditions (**FIG 8B**), but the effect was small and the physiological relevance of this result is unclear. Based on these results, we conclude that neither RclB nor RclC is likely to function by regulating the bulk synthesis or degradation of any of the major OMPs, but this does not preclude the possibility that they interact with one or more of those proteins to regulate their functions or the ability of HOSCN to diffuse through their pores. We are currently pursuing experiments to test this possibility.

**FIG 8.**
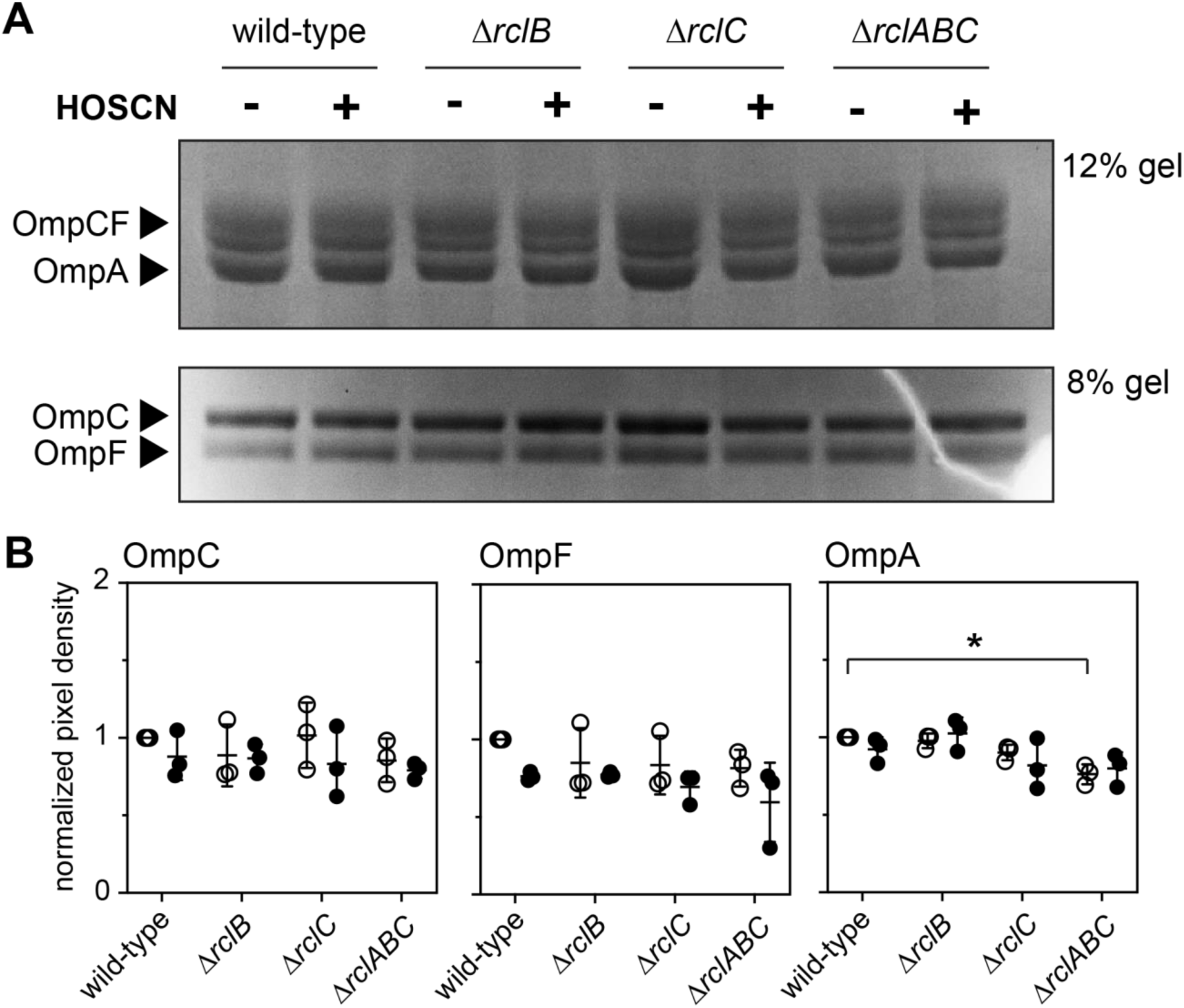
HOSCN stress does not have major impacts on the abundance of OmpC, OmpF, or OmpA in *E. coli*. *E. coli* MG1655 wild-type, MJG0047 (MG1655 Δ*rclB*), MJG0013 (MG1655 Δ*rclC*::*cat*^+^), and MJG0901 (MG1655 Δ*rclABC*) were grown at 37°C with shaking to A_600_ = 0.3 in M9 minimal medium, then 200 µM HOSCN was added to treatment samples. Cultures were incubated at 37°C with shaking to A_600_ = 1.2 – 1.3, then outer membrane protein fractions were isolated and separated on (**A**) 12% or 8% acrylamide Bolt^TM^ SDS-PAGE gels (Invitrogen) and visualized by Coomassie Blue staining. Complete gel images from triplicates of this experiment are shown in **SUPPLEMENTAL FIG S3**. (**B**) The indicated protein band intensities were measured with ImageJ (88) and normalized to their intensities in untreated wild-type *E. coli* for each experiment. Statistical significance calculated in Prism GraphPad by two-way ANOVA: * = P-value < 0.05. Full ANOVA results in **SUPPLEMENTAL TABLE S1**.

## CONCLUSIONS

### A connection between HOSCN, the *rcl* operon, and envelope permeability

In this study, we have characterized the overall transcriptomic response of *E. coli* MG1655 to treatment by HOSCN for the first time. We observed that *E. coli* has a more specific response to HOSCN than what has been shown in other immune-derived oxidants, with many of the typical oxidative stress responses being only partially upregulated (**FIG 1**). When any of the *rcl* genes were mutated, *E. coli* demonstrated both greater sensitivity (**FIG 2**) and a greater transcriptional response to HOSCN (**FIGS 3**, **4**), although surprisingly, the transcriptional response to HOSCN was mostly highly comparable in all knockout strains, and not dependent on one *rcl* gene over the other. Transcriptional signatures and protein homology (**FIGS 4**, **6**, and **7**) lead us to suggest a model (**FIG 9**) in which RclA, RclB, and RclC act as a coordinated HOSCN defense system, with RclB and RclC acting (likely together, based on the similarity of the phenotypes of Δ*rclB* and Δ*rclC* mutants)(**FIGS 2**, **4F**) to prevent penetration of HOSCN into the *E. coli* cytoplasm. We hypothesize based on structural and sequence homology to other proteins that RclB may do this by interaction with OMPs (**FIGS 6**, **7**, and **8**), but the precise molecular mechanism by which it might do so and what role the inner membrane protein RclC may play in this process remain to be determined.

**FIG 9.**
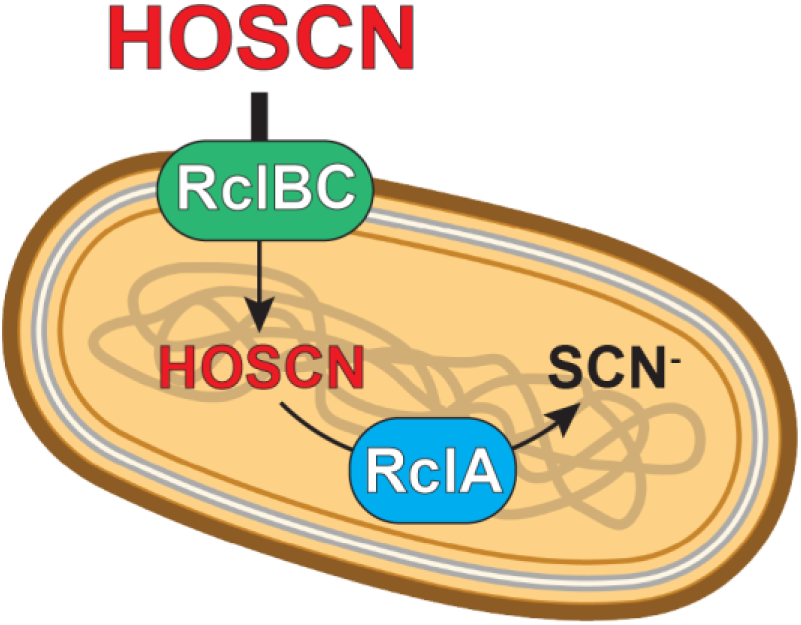
Model for the roles of RclABC in HOSCN stress resistance in *E. coli*. Based on the results presented here, we propose a model in which RclB and RclC work together to limit the permeability of the *E. coli* cell envelope to HOSCN, possibly by regulating the function of the major outer membrane porins. Once HOSCN penetrates the cytoplasm, it is degraded to non-toxic SCN^-^ by the HOSCN reductase RclA (20).

## MATERIALS AND METHODS

### Bacterial strain construction

All bacterial strains used in this study can be found in Table 1. The *E. coli* Δ*ompC* and Δ*ompF* strains were generated by transducing the Δ*ompC::kan^+^* and Δ*ompF::kan^+^* alleles from the Keio collection (78) into MG1655 (79) using P1*vir* phage (80). This generated strains MJG2366 and MJG2367 respectively. The kanamycin resistance cassette in both strains was resolved (81) to yield strains Δ*ompC* (MJG2411) and Δ*ompF* (MJG2412). The double knockout strain MJG2429 (Δ*ompCF*) was generated by transducing the Δ*ompF::kan^+^* allele into MJG2411 using P1*vir*.

**TABLE 1.** Strains and plasmids used in this study. Unless otherwise indicated, strains and plasmids were generated in the course of this work. Abbreviations: Cm^R^, chloramphenicol resistance; Kn^R^, kanamycin resistance.

| Strain | Marker | Genotype | Reference |
| --- | --- | --- | --- |
| MG1655 |  | F<sup>-</sup>, λ<sup>-</sup>, *rph-1 ilvG<sup>-</sup> rfb-50* | (79) |
| MJG0013 | Cm<sup>R</sup> | MG1655 $\Delta rclC745::cat^{+}$ | (33) |
| MJG0047 | | MG1655 $\Delta rclB746$ | (33) |
| MJG1958 | | MG1655 $\Delta rclA747$ | (20) |
| MJG0901 | | MG1655 $\Delta rclABC$ | (32) |
| MJG2366 | Kn<sup>R</sup> | MG1655 $\Delta ompC768::kan^{+}$ | |
| MJG2367 | Kn<sup>R</sup> | MG1655 $\Delta ompF746::kan^{+}$ | |
| MJG2411 | | MG1655 $\Delta ompC768$ | |
| MJG2412 | | MG1655 $\Delta ompF746$ | |
| MJG2429 | Kn<sup>R</sup> | MG1655 $\Delta ompC768 \Delta ompF746::kan^{+}$ | |

### RNA sequencing

For RNA extraction, single colonies of *E. coli* were inoculated into 5mL of LB and grown overnight at 37°C with shaking. The next day, *E. coli* was subcultured into M9 minimal media containing 100 μM FeCl_3_ and grown to mid log phase (absorbance at 600 nm = 0.4-0.5). Cultures were treated with 600 μM HOSCN for 15 minutes, at which point they were harvested, and RNA was extracted using the Qiagen RNeasy PowerMicrobiome Kit. mRNA sequencing was carried out by SeqCenter (Pittsburgh, PA). Samples were DNAse treated with Invitrogen DNAse (RNAse free). Library preparation was performed using Illumina’s Stranded Total RNA Prep Ligation with Ribo-Zero Plus kit and 10 bp unique dual indices (UDI). Sequencing was done on a NovaSeq X Plus, producing paired end 150 bp reads. Demultiplexing, quality control, and adapter trimming was performed with bcl-convert (v4.1.5). Quality control and adapter trimming was performed with bcl-convert. Read mapping was performed with HISAT2 (82). Read quantification was performed using Subread’s featureCounts functionality (83). Read counts loaded into R were normalized using edgeR’s Trimmed Mean of M values (TMM) algorithm. Subsequent values were then converted to counts per million (CPM). Differential expression analysis was performed using edgeR’s glmQLFTest.

### Data availability

The raw data for the transcriptomic experiment described above is available in the NIH Sequence Read Archive (accession number PRJNA989513), and a human-readable version (with CPMs converted to CPM+1 to allow for calculation of log_2_-fold change in expression for all genes, including those with no detectable reads under some conditions) is available as **SUPPLEMENTAL DATA SET 1**.

### Measuring bacterial growth under HOSCN stress

Single colonies of *E. coli* were inoculated in M9 minimal media (84) containing 0.2% glucose and 100 μM FeCl_3_ and grown overnight at 37°C with shaking. The next day, cultures were harvested and normalized to OD_600_=0.05 in a 96-well plate with M9 minimal media containing the indicated concentrations of HOSCN. The plate was covered with a BreatheEasy plate film (Andwin Scientific) and placed in a Tecan Sunrise plate reader where the absorbance was measured at 600 nm every 15 minutes for 24 hours, with shaking in between each measurement. HOSCN was made and quantified fresh the day of the experiment as previously described (20). Due to the highly reactive nature of HOSCN, the exact period of growth inhibition caused by a given dose of HOSCN varied from day to day, so representative growth curve data are shown in **FIGS 2** and **5**, with data from additional independent experimental replicates shown in **SUPPLEMENTAL FIGS S1** and **S2**, respectively to illustrate the consistency in relative sensitivity of mutant strains to the wild-type and to one another.

### Structural homology

Structural homology of RclB was investigated with the COFACTOR tool, using the default parameters (76). The pdb file of the predicted structure of RclB was obtained from AlphaFold (85, 86). The ribbon structure images were produced using Pymol (The PyMOL Molecular Graphics System, Version 3.0 Schrödinger, LLC.).

### Cell collection for isolation of outer membrane proteins

Single colonies of *E. coli* were inoculated into 5 mL of LB and grown overnight at 37°C with shaking. The next day, 500 μL of overnight was subcultured into 25 mL of M9 minimal growth media and grown to early log phase (absorbance at 600 nm = 0.3), at which point treatment samples were dosed with HOSCN to a final concentration of 200 μM, then continued to grow until the optical density reached 1.2-1.3. Cell pellets were spun down and stored at -80°C until needed.

### Outer membrane protein preparation

Outer membrane fractions were prepared by a modification of a previously published method (87). Frozen cell pellets were thawed at room temperature, then resuspended in PBS. Cultures were sonicated to break open cells for 2 minutes, 5 seconds on 5 seconds off, at 50% amplification, 50 V using a Model 120 Sonic Dismembrator (Fisher). Samples were then centrifuged at 1400 x g for 10 minutes to collect cell debris. Supernatants were centrifuged at 85,000 x g for 40 minutes in a Beckman-Coulter Optima XPN-80 ultracentrifuge, then solubilized in a solution of 2% Triton-X 100 in Tris-HCl (pH 7.5) for 30 minutes at 37°C. Outer membranes were then collected by ultracentrifugation at 85,000 x g for 40 minutes.

Proteins were visualized by running on 8% or 12% Invitrogen Bolt gels with MOPS buffer and staining with Coomassie Blue, and band intensity was quantified with ImageJ (88).

## REFERENCES

1. Chaplin DD. 2010. Overview of the immune response. J Allergy Clin Immunol 125:S3–23.

2. Martens EC, Neumann M, Desai MS. 2018. Interactions of commensal and pathogenic microorganisms with the intestinal mucosal barrier. Nat Rev Microbiol 16:457–470.

3. Nathan C, Cunningham-Bussel A. 2013. Beyond oxidative stress: an immunologist’s guide to reactive oxygen species. Nat Rev Immunol 13:349–61.

4. Winterbourn CC, Kettle AJ, Hampton MB. 2016. Reactive Oxygen Species and Neutrophil Function. Annu Rev Biochem 85:765–92.

5. Barrett TJ, Hawkins CL. 2012. Hypothiocyanous acid: benign or deadly? Chem Res Toxicol 25:263–73.

6. Meredith JD, Gray MJ. 2023. Hypothiocyanite and host-microbe interactions. Mol Microbiol 119:302–311.

7. Day BJ. 2019. The science of licking your wounds: Function of oxidants in the innate immune system. Biochem Pharmacol 163:451–457.

8. Ulfig A, Leichert LI. 2021. The effects of neutrophil-generated hypochlorous acid and other hypohalous acids on host and pathogens. Cell Mol Life Sci 78:385–414.

9. Reiter B, Harnulv G. 1984. Lactoperoxidase Antibacterial System: Natural Occurrence, Biological Functions and Practical Applications. J Food Prot 47:724–732.

10. Arnhold J, Malle E. 2022. Halogenation Activity of Mammalian Heme Peroxidases. Antioxidants (Basel) 11.

11. Ramirez GA, Yacoub MR, Ripa M, Mannina D, Cariddi A, Saporiti N, Ciceri F, Castagna A, Colombo G, Dagna L. 2018. Eosinophils from Physiology to Disease: A Comprehensive Review. Biomed Res Int 2018:9095275.

12. Pattison DI, Davies MJ, Hawkins CL. 2012. Reactions and reactivity of myeloperoxidase-derived oxidants: differential biological effects of hypochlorous and hypothiocyanous acids. Free Radic Res 46:975–95.

13. Ashby MT. 2012. Chapter 8 - Hypothiocyanite, p 263–303. In Eldik Rv, Ivanović-Burmazović I (ed), Advances in Inorganic Chemistry: Inorganic/Bioinorganic Reaction Mechanisms, vol 64. Elsevier Inc.

14. Skaff O, Pattison DI, Davies MJ. 2009. Hypothiocyanous acid reactivity with low-molecular-mass and protein thiols: absolute rate constants and assessment of biological relevance. Biochem J 422:111–7.

15. Imlay JA. 2008. Cellular defenses against superoxide and hydrogen peroxide. Annu Rev Biochem 77:755–76.

16. Gray MJ, Wholey WY, Jakob U. 2013. Bacterial responses to reactive chlorine species. Annu Rev Microbiol 67:141–60.

17. Yesilkaya H, Andisi VF, Andrew PW, Bijlsma JJ. 2013. Streptococcus pneumoniae and reactive oxygen species: an unusual approach to living with radicals. Trends Microbiol 21:187–95.

18. Hamon CB, Klebanoff SJ. 1973. A peroxidase-mediated, streptococcus mitis-dependent antimicrobial system in saliva. J Exp Med 137:438–50.

19. Oram JD, Reiter B. 1966. The inhibition of streptococci by lactoperoxidase, thiocyanate and hydrogen peroxide. The effect of the inhibitory system on susceptible and resistant strains of group N streptococci. Biochem J 100:373–81.

20. Meredith JD, Chapman I, Ulrich K, Sebastian C, Stull F, Gray MJ. 2022. Escherichia coli RclA is a highly active hypothiocyanite reductase. Proc Natl Acad Sci U S A 119:e2119368119.

21. Shearer HL, Pace PE, Paton JC, Hampton MB, Dickerhof N. 2022. A newly identified flavoprotein disulfide reductase Har protects Streptococcus pneumoniae against hypothiocyanous acid. J Biol Chem doi:10.1016/j.jbc.2022.102359:102359.

22. Shearer HL, Loi VV, Weiland P, Bange G, Altegoer F, Hampton MB, Antelmann H, Dickerhof N. 2023. MerA functions as a hypothiocyanous acid reductase and defense mechanism in Staphylococcus aureus. Mol Microbiol 119:456–470.

23. Chandler JD, Nichols DP, Nick JA, Hondal RJ, Day BJ. 2013. Selective metabolism of hypothiocyanous acid by mammalian thioredoxin reductase promotes lung innate immunity and antioxidant defense. J Biol Chem 288:18421–8.

24. Imlay JA. 2015. Transcription Factors That Defend Bacteria Against Reactive Oxygen Species. Annu Rev Microbiol 69:93–108.

25. Gray MJ. 2024. The role of metals in hypothiocyanite resistance in Escherichia coli. J Bacteriol 206:e0009824.

26. Shearer HL, Currie MJ, Agnew HN, Trappetti C, Stull F, Pace PE, Paton JC, Dobson RCJ, Dickerhof N. 2024. Hypothiocyanous acid reductase is critical for host colonization and infection by Streptococcus pneumoniae. J Biol Chem 300:107282.

27. Shearer HL, Paton JC, Hampton MB, Dickerhof N. 2022. Glutathione utilization protects Streptococcus pneumoniae against lactoperoxidase-derived hypothiocyanous acid. Free Radic Biol Med 179:24–33.

28. Farrant KV, Spiga L, Davies JC, Williams HD. 2020. Response of Pseudomonas aeruginosa to the Innate Immune System-Derived Oxidants Hypochlorous Acid and Hypothiocyanous Acid. J Bacteriol 203.

29. Groitl B, Dahl JU, Schroeder JW, Jakob U. 2017. Pseudomonas aeruginosa defense systems against microbicidal oxidants. Mol Microbiol 106:335–350.

30. Derke RM, Barron AJ, Billiot CE, Chaple IF, Lapi SE, Broderick NA, Gray MJ. 2020. The Cu(II) Reductase RclA Protects Escherichia coli against the Combination of Hypochlorous Acid and Intracellular Copper. mBio 11.

31. Shearer HL, Pace PE, Smith LM, Fineran PC, Matthews AJ, Camilli A, Dickerhof N, Hampton MB. 2023. Identification of Streptococcus pneumoniae genes associated with hypothiocyanous acid tolerance through genome-wide screening. J Bacteriol 205:e0020823.

32. Konigstorfer A, Ashby LV, Bollar GE, Billiot CE, Gray MJ, Jakob U, Hampton MB, Winterbourn CC. 2021. Induction of the reactive chlorine-responsive transcription factor RclR in Escherichia coli following ingestion by neutrophils. Pathog Dis 79.

33. Parker BW, Schwessinger EA, Jakob U, Gray MJ. 2013. The RclR protein is a reactive chlorine-specific transcription factor in Escherichia coli. J Biol Chem 288:32574–32584.

34. Sung YC, Fuchs JA. 1988. Characterization of the cyn operon in Escherichia coli K12. J Biol Chem 263:14769–75.

35. Sung YC, Fuchs JA. 1992. The Escherichia coli K-12 cyn operon is positively regulated by a member of the lysR family. J Bacteriol 174:3645–50.

36. Hensley MP, Gunasekera TS, Easton JA, Sigdel TK, Sugarbaker SA, Klingbeil L, Breece RM, Tierney DL, Crowder MW. 2012. Characterization of Zn(II)-responsive ribosomal proteins YkgM and L31 in E. coli. J Inorg Biochem 111:164–72.

37. Graham AI, Hunt S, Stokes SL, Bramall N, Bunch J, Cox AG, McLeod CW, Poole RK. 2009. Severe zinc depletion of Escherichia coli: roles for high affinity zinc binding by ZinT, zinc transport and zinc-independent proteins. J Biol Chem 284:18377–89.

38. Petrarca P, Ammendola S, Pasquali P, Battistoni A. 2010. The Zur-regulated ZinT protein is an auxiliary component of the high-affinity ZnuABC zinc transporter that facilitates metal recruitment during severe zinc shortage. J Bacteriol 192:1553–64.

39. Patzer SI, Hantke K. 2000. The zinc-responsive regulator Zur and its control of the znu gene cluster encoding the ZnuABC zinc uptake system in Escherichia coli. J Biol Chem 275:24321–32.

40. Maret W, Li Y. 2009. Coordination dynamics of zinc in proteins. Chem Rev 109:4682–707.

41. Chubiz LM. 2023. The Mar, Sox, and Rob Systems. EcoSal Plus 11:eesp00102022.

42. Moreau PL. 2007. The lysine decarboxylase CadA protects Escherichia coli starved of phosphate against fermentation acids. J Bacteriol 189:2249–61.

43. Al-Fageeh MB, Smales CM. 2006. Control and regulation of the cellular responses to cold shock: the responses in yeast and mammalian systems. Biochem J 397:247–59.

44. Zhang Y, Gross CA. 2021. Cold Shock Response in Bacteria. Annu Rev Genet 55:377–400.

45. Kittleson JT, Loftin IR, Hausrath AC, Engelhardt KP, Rensing C, McEvoy MM. 2006. Periplasmic metal-resistance protein CusF exhibits high affinity and specificity for both CuI and AgI. Biochemistry 45:11096–102.

46. Gustavsson N, Diez A, Nystrom T. 2002. The universal stress protein paralogues of Escherichia coli are co-ordinately regulated and co-operate in the defence against DNA damage. Mol Microbiol 43:107–17.

47. Gray MJ, Wholey WY, Parker BW, Kim M, Jakob U. 2013. NemR is a bleach-sensing transcription factor. J Biol Chem 288:13789–98.

48. Gebendorfer KM, Drazic A, Le Y, Gundlach J, Bepperling A, Kastenmuller A, Ganzinger KA, Braun N, Franzmann TM, Winter J. 2012. Identification of a hypochlorite-specific transcription factor from Escherichia coli. J Biol Chem 287:6892–903.

49. Sultana S, Foti A, Dahl JU. 2020. Bacterial Defense Systems against the Neutrophilic Oxidant Hypochlorous Acid. Infect Immun 88.

50. Mermod M, Magnani D, Solioz M, Stoyanov JV. 2012. The copper-inducible ComR (YcfQ) repressor regulates expression of ComC (YcfR), which affects copper permeability of the outer membrane of Escherichia coli. Biometals 25:33–43.

51. Zhang XS, Garcia-Contreras R, Wood TK. 2007. YcfR (BhsA) influences Escherichia coli biofilm formation through stress response and surface hydrophobicity. J Bacteriol 189:3051–62.

52. Karp PD, Paley S, Caspi R, Kothari A, Krummenacker M, Midford PE, Moore LR, Subhraveti P, Gama-Castro S, Tierrafria VH, Lara P, Muniz-Rascado L, Bonavides-Martinez C, Santos-Zavaleta A, Mackie A, Sun G, Ahn-Horst TA, Choi H, Covert MW, Collado-Vides J, Paulsen I. 2023. The EcoCyc Database (2023). EcoSal Plus 11:eesp00022023.

53. Buchbinder JL, Stephenson RC, Dresser MJ, Pitera JW, Scanlan TS, Fletterick RJ. 1998. Biochemical characterization and crystallographic structure of an Escherichia coli protein from the phosphotriesterase gene family. Biochemistry 37:5096–106.

54. Ma Q, Yang Z, Pu M, Peti W, Wood TK. 2011. Engineering a novel c-di-GMP-binding protein for biofilm dispersal. Environ Microbiol 13:631–42.

55. Das U, Shuman S. 2013. 2’-Phosphate cyclase activity of RtcA: a potential rationale for the operon organization of RtcA with an RNA repair ligase RtcB in Escherichia coli and other bacterial taxa. RNA 19:1355–62.

56. Blum M, Andreeva A, Florentino LC, Chuguransky SR, Grego T, Hobbs E, Pinto BL, Orr A, Paysan-Lafosse T, Ponamareva I, Salazar GA, Bordin N, Bork P, Bridge A, Colwell L, Gough J, Haft DH, Letunic I, Llinares-Lopez F, Marchler-Bauer A, Meng-Papaxanthos L, Mi H, Natale DA, Orengo CA, Pandurangan AP, Piovesan D, Rivoire C, Sigrist CJA, Thanki N, Thibaud-Nissen F, Thomas PD, Tosatto SCE, Wu CH, Bateman A. 2024. InterPro: the protein sequence classification resource in 2025. Nucleic Acids Res doi:10.1093/nar/gkae1082.

57. Eletsky A, Michalska K, Houliston S, Zhang Q, Daily MD, Xu X, Cui H, Yee A, Lemak A, Wu B, Garcia M, Burnet MC, Meyer KM, Aryal UK, Sanchez O, Ansong C, Xiao R, Acton TB, Adkins JN, Montelione GT, Joachimiak A, Arrowsmith CH, Savchenko A, Szyperski T, Cort JR. 2014. Structural and functional characterization of DUF1471 domains of Salmonella proteins SrfN, YdgH/SssB, and YahO. PLoS One 9:e101787.

58. Crompton ME, Gaessler LF, Tawiah PO, Polzer L, Camfield SK, Jacobson GD, Naudszus MK, Johnson C, Meurer K, Bennis M, Roseberry B, Sultana S, Dahl JU. 2023. Expression of RcrB confers resistance to hypochlorous acid in uropathogenic Escherichia coli. J Bacteriol 205:e0006423.

59. Kobylka J, Kuth MS, Muller RT, Geertsma ER, Pos KM. 2020. AcrB: a mean, keen, drug efflux machine. Ann N Y Acad Sci 1459:38–68.

60. Patzer SI, Hantke K. 1998. The ZnuABC high-affinity zinc uptake system and its regulator Zur in Escherichia coli. Mol Microbiol 28:1199–210.

61. Fernandes AP, Holmgren A. 2004. Glutaredoxins: glutathione-dependent redox enzymes with functions far beyond a simple thioredoxin backup system. Antioxid Redox Signal 6:63–74.

62. Nagy P, Jameson GN, Winterbourn CC. 2009. Kinetics and mechanisms of the reaction of hypothiocyanous acid with 5-thio-2-nitrobenzoic acid and reduced glutathione. Chem Res Toxicol 22:1833–40.

63. Loi VV, Busche T, Schnaufer F, Kalinowski J, Antelmann H. 2023. The neutrophil oxidant hypothiocyanous acid causes a thiol-specific stress response and an oxidative shift of the bacillithiol redox potential in Staphylococcus aureus. Microbiol Spectr 11:e0325223.

64. Perham RN. 1987. Glutathione reductase from Escherichia coli: mutation, cloning and sequence analysis of the gene. Biochem Soc Trans 15:730–3.

65. Kobayashi K, Fujikawa M, Kozawa T. 2014. Oxidative stress sensing by the iron-sulfur cluster in the transcription factor, SoxR. J Inorg Biochem 133:87–91.

66. Gray MJ, Li Y, Leichert LI, Xu Z, Jakob U. 2015. Does the Transcription Factor NemR Use a Regulatory Sulfenamide Bond to Sense Bleach? Antioxid Redox Signal 23:747–54.

67. Weissbach H, Brot N. 1991. Regulation of methionine synthesis in Escherichia coli. Mol Microbiol 5:1593–7.

68. Lugtenberg B, Van Alphen L. 1983. Molecular architecture and functioning of the outer membrane of Escherichia coli and other gram-negative bacteria. Biochim Biophys Acta 737:51–115.

69. De la Cruz MA, Calva E. 2010. The complexities of porin genetic regulation. J Mol Microbiol Biotechnol 18:24–36.

70. Nikaido H. 2003. Molecular basis of bacterial outer membrane permeability revisited. Microbiol Mol Biol Rev 67:593–656.

71. De Spiegeleer P, Sermon J, Vanoirbeek K, Aertsen A, Michiels CW. 2005. Role of porins in sensitivity of Escherichia coli to antibacterial activity of the lactoperoxidase enzyme system. Appl Environ Microbiol 71:3512–8.

72. Kim S, Song I, Eom G, Kim S. 2018. A Small Periplasmic Protein with a Hydrophobic C-Terminal Residue Enhances DegP Proteolysis as a Suicide Activator. J Bacteriol 200.

73. Lee J, Hiibel SR, Reardon KF, Wood TK. 2010. Identification of stress-related proteins in Escherichia coli using the pollutant cis-dichloroethylene. J Appl Microbiol 108:2088–102.

74. Zhang XS, Garcia-Contreras R, Wood TK. 2008. Escherichia coli transcription factor YncC (McbR) regulates colanic acid and biofilm formation by repressing expression of periplasmic protein YbiM (McbA). ISME J 2:615–31.

75. Sargentini NJ, Gularte NP, Hudman DA. 2016. Screen for genes involved in radiation survival of Escherichia coli and construction of a reference database. Mutat Res 793–794:1-14.

76. Zhang C, Freddolino PL, Zhang Y. 2017. COFACTOR: improved protein function prediction by combining structure, sequence and protein-protein interaction information. Nucleic Acids Res 45:W291–W299.

77. Cho SH, Dekoninck K, Collet JF. 2023. Envelope-Stress Sensing Mechanism of Rcs and Cpx Signaling Pathways in Gram-Negative Bacteria. J Microbiol 61:317–329.

78. Baba T, Ara T, Hasegawa M, Takai Y, Okumura Y, Baba M, Datsenko KA, Tomita M, Wanner BL, Mori H. 2006. Construction of Escherichia coli K-12 in-frame, single-gene knockout mutants: the Keio collection. Mol Syst Biol 2:2006 0008.

79. Blattner FR, Plunkett G, 3rd, Bloch CA, Perna NT, Burland V, Riley M, Collado-Vides J, Glasner JD, Rode CK, Mayhew GF, Gregor J, Davis NW, Kirkpatrick HA, Goeden MA, Rose DJ, Mau B, Shao Y. 1997. The complete genome sequence of Escherichia coli K-12. Science 277:1453–62.

80. Silhavy TJ, Berman ML, Enquist LW. 1984. Experiments with gene fusions. Cold Springs Harbor Laboratory, Cold Springs Harbor, NY.

81. Datsenko KA, Wanner BL. 2000. One-step inactivation of chromosomal genes in Escherichia coli K-12 using PCR products. Proc Natl Acad Sci U S A 97:6640–5.

82. Kim D, Paggi JM, Park C, Bennett C, Salzberg SL. 2019. Graph-based genome alignment and genotyping with HISAT2 and HISAT-genotype. Nat Biotechnol 37:907–915.

83. Liao Y, Smyth GK, Shi W. 2014. featureCounts: an efficient general purpose program for assigning sequence reads to genomic features. Bioinformatics 30:923–30.

84. Sambrook J, E. F. Fritsch and T. Maniatis. 1989. Molecular cloning: a laboratory manual, 2nd ed. Cold Spring Harbor Laboratory, Cold Spring Harbor, N.Y.

85. Varadi M, Anyango S, Deshpande M, Nair S, Natassia C, Yordanova G, Yuan D, Stroe O, Wood G, Laydon A, Zidek A, Green T, Tunyasuvunakool K, Petersen S, Jumper J, Clancy E, Green R, Vora A, Lutfi M, Figurnov M, Cowie A, Hobbs N, Kohli P, Kleywegt G, Birney E, Hassabis D, Velankar S. 2022. AlphaFold Protein Structure Database: massively expanding the structural coverage of protein-sequence space with high-accuracy models. Nucleic Acids Res 50:D439–D444.

86. Jumper J, Evans R, Pritzel A, Green T, Figurnov M, Ronneberger O, Tunyasuvunakool K, Bates R, Zidek A, Potapenko A, Bridgland A, Meyer C, Kohl SAA, Ballard AJ, Cowie A, Romera-Paredes B, Nikolov S, Jain R, Adler J, Back T, Petersen S, Reiman D, Clancy E, Zielinski M, Steinegger M, Pacholska M, Berghammer T, Bodenstein S, Silver D, Vinyals O, Senior AW, Kavukcuoglu K, Kohli P, Hassabis D. 2021. Highly accurate protein structure prediction with AlphaFold. Nature 596:583–589.

87. Lin X, Wang C, Guo C, Tian Y, Li H, Peng X. 2012. Differential regulation of OmpC and OmpF by AtpB in Escherichia coli exposed to nalidixic acid and chlortetracycline. J Proteomics 75:5898–910.

88. Schneider CA, Rasband WS, Eliceiri KW. 2012. NIH Image to ImageJ: 25 years of image analysis. Nat Methods 9:671–5.

89. Madeira F, Madhusoodanan N, Lee J, Eusebi A, Niewielska A, Tivey ARN, Lopez R, Butcher S. 2024. The EMBL-EBI Job Dispatcher sequence analysis tools framework in 2024. Nucleic Acids Res 52:W521–W525.

90. Leverrier P, Declercq JP, Denoncin K, Vertommen D, Hiniker A, Cho SH, Collet JF. 2011. Crystal structure of the outer membrane protein RcsF, a new substrate for the periplasmic protein-disulfide isomerase DsbC. J Biol Chem 286:16734–42.

